# VirClust – a tool for hierarchical clustering, core gene detection and annotation of (prokaryotic) viruses

**DOI:** 10.1101/2021.06.14.448304

**Authors:** Cristina Moraru

## Abstract

Recent years have seen major changes in the classification criteria and taxonomy of viruses. The current classification scheme, also called “megataxonomy of viruses”, recognizes five different viral realms, defined based on the presence of viral hallmark genes. Within the realms, viruses are classified into hierarchical taxons, ideally defined by their shared genes. Therefore, there is currently a need for virus classification tools based on such shared genes / proteins. Here, VirClust is presented – a novel tool capable of performing i) hierarchical clustering of viruses based on intergenomic distances calculated from their protein cluster content, ii) identification of core proteins and iii) annotation of viral proteins. VirClust groups proteins into clusters both based on BLASTP sequence similarity, which identifies more related proteins, and also based on hidden markow models (HMM), which identifies more distantly related proteins. Furthermore, VirClust provides an integrated visualization of the hierarchical clustering tree and of the distribution of the protein content, which allows the identification of the genomic features responsible for the respective clustering. By using different intergenomic distances, the hierarchical trees produced by VirClust can be split into viral genome clusters of different taxonomic ranks. VirClust is freely available, as web-service (virclust.icbm.de) and stand-alone tool.

## Introduction

Viral classification and taxonomy has recently undergone major changes. The Baltimore classification scheme, based solely on the viral nucleic acid type has been replaced by a viral megataxonomy, based on viral genome features, including shared genes (proteins) (Koonin *et al*. 2020). The traditional five-rank structure of viral taxonomy was replaced by a 15-rank classification hierarchy, similar to the Linean taxonomy (Gorbalenya *et al*. 2020). As catalyst for these changes served the unparalleled insights into virus genome organization and evolution facilitated by the advent of genome sequencing.

In contrast to cellular organisms, which all share a common ancestor and have preserved a number of universal genes, viruses share no universal gene and likely have multiple points of origin (Koonin *et al*. 2006; Krupovic and Koonin 2017; Krupovic *et al*. 2019; Kazlauskas *et al*. 2019). Therefore, traditional phylogenetic methods, in which phylogenetic trees are constructed based on multiple alignments of homologous genes (proteins), cannot be applied to viruses as a whole. Gene (protein) sharing networks have been used to explore how viruses are related with each other (Iranzo *et al*. 2016) and resulted in the definition of viral hallmark genes (VHGs), which represent genes broadly found in diverse virus groups, but not universally present. Based on the presence of such VHGs, five viral realms have been defined to date: *Adnaviria, Riboviria, Duplodnaviria* and *Varidnaviria* (Koonin *et al*. 2020; Krupovic *et al*. 2021). Prokaryotic viruses, infecting bacteria and archaea, are spread through the five realms, with the known majority belonging to the order *Caudovirales*, within *Duplodnaviria*. Many others, cultivated and uncultivated, are yet unassigned to any realm, awainting further evidence to classify them into an already existing realm or to a brand new one (Koonin *et al*. 2020).

Inside the realms, viruses are further organized into hierarchical taxons, from kingdom to species, similar to the cellular world (Gorbalenya *et al*. 2020). At lower-rank levels – species and genus –, the classification is based on intergenomic nucleic acid identities, as calculated for example with VIRIDIC (Moraru *et al*. 2020). The Bacterial and Archaeal Viruses Subcommittee of International Committtee on Taxonomy of Viruses (ICTV) recommends a 95% and 70% identity threshold for the species and genus level, respectively. The monophyly of genus level clusters should be evaluated using core/signature gene phylogeny. The classification criteria for the intermediary-level ranks, as for example family and order, are currently being defined. They should be based on whole viral proteomes and should take into account shared orthologous proteins (Turner *et al*. 2021). Further on, at the highest-level rank, the realm, viruses should be classified based on the presence of specific VHGs, which can be recognized through protein clustering and annotation methods.

There are a number of whole proteome-based virus classification tools, used also for the delineation of intermediary ranks. They can be classified in tools based on i) whole proteome similarity, as for example ViPTree (Nishimura *et al*. 2017) and VICTOR (Meier-Kolthoff and Göker 2017), ii) on protein profile hidden Markov models (PPHMM) and genomic organization models (GOM), as implemented in GRAViTy (Aiewsakun and Simmonds 2018; Aiewsakun *et al*. 2018) and iii) on shared protein clusters, as implemented in vConTACT (Bolduc *et al*. 2017; Bin Jang *et al*. 2019). VICTOR and VipTree calculate pairwise intergenomic distances based on protein-protein BLAST comparisons of the whole viral proteomes in a given dataset, and use them for clustering hierarchically the respective viruses. GRAViTy uses concatenated proteins of the query viruses to search against pre-calculated databases of viral PPHMMs and GOMs and then computes for each query virus a PPHMM and GOM signature. These signatures contain information about the degree of similarity between the query and the databases and are used to calculate intergenomic pairwise distances, followed by hierarchical clustering of the viruses. Finaly, vConTACT computes for the given dataset of viral genomes (including or not a reference database) all protein clusters, based on BLASTP comparisons. Then, it uses the absence/presence of protein clusters to calculate intergenomic similarities between viruses, which are further used to construct a viral genome monopartite network. This method produces single-level viral clusters, potentially of the genus or family rank. The main disadvantages of these tools is that they either don’t identify the genomic features (proteins) contributing to the clustering of the viruses (ViPTree, VICTOR, GRAViTy), or they don’t produce hierarchical clusters (vConTACT).

VirClust (<u>v</u>irus <u>c</u>lusterer), presented here, is meant to complement the existing viral classification tools, by bringing to the table the following: i) calculation of intergenomic distances based on the presence/absence of protein clusters, clusters determined by BLASTP or HMM profile comparisions; ii) hierarchical clustering of the viral genomes based on the respective intergenomic distances; iii) integrated visualization of the viral clusters and their protein cluster content; iv) calculation of core protein clusters; and v) protein annotation based on a state of the art colection of sequence databases. VirClust is available both as a web-service (virclust.icbm.de) and as a stand-alone command-line tool.

## Methods

### VirClust – development and workflow

VirClust was developed in the R v3.5 (R Core Team 2018) programming language. The web interface was developed under the Shiny web application framework (https://cran.r-project.org/web/packages/shiny/index.html, RStudio, MA, USA). The stand-alone tool for Linux was wrapped in a container using the Singularity v. 3.5.2 software (https://sylabs.io/, Sylabs.io, CA, USA). The stand-alone version can be deployed on any systems running the Singularity software.

A complete VirClust workflow has 8 steps, each with its own individual outputs. These outputs, referred from here on as “usable outputs” (see Figure 1), can be retrieved by the user either by download from the webpage (when using the VirClust web-service) or directly from the disk space, when using the VirClust standalone version. The user can chose which steps to run. If a certain step depends on the output from a previous step, it will be automatically activated by VirClust (for example, if the user choses step4b, which depends on step3b, step3b will be performed as well). The “continue” option allows to run new steps (or to re-run some steps) in the workflow at a later time point.

**Figure 1:**
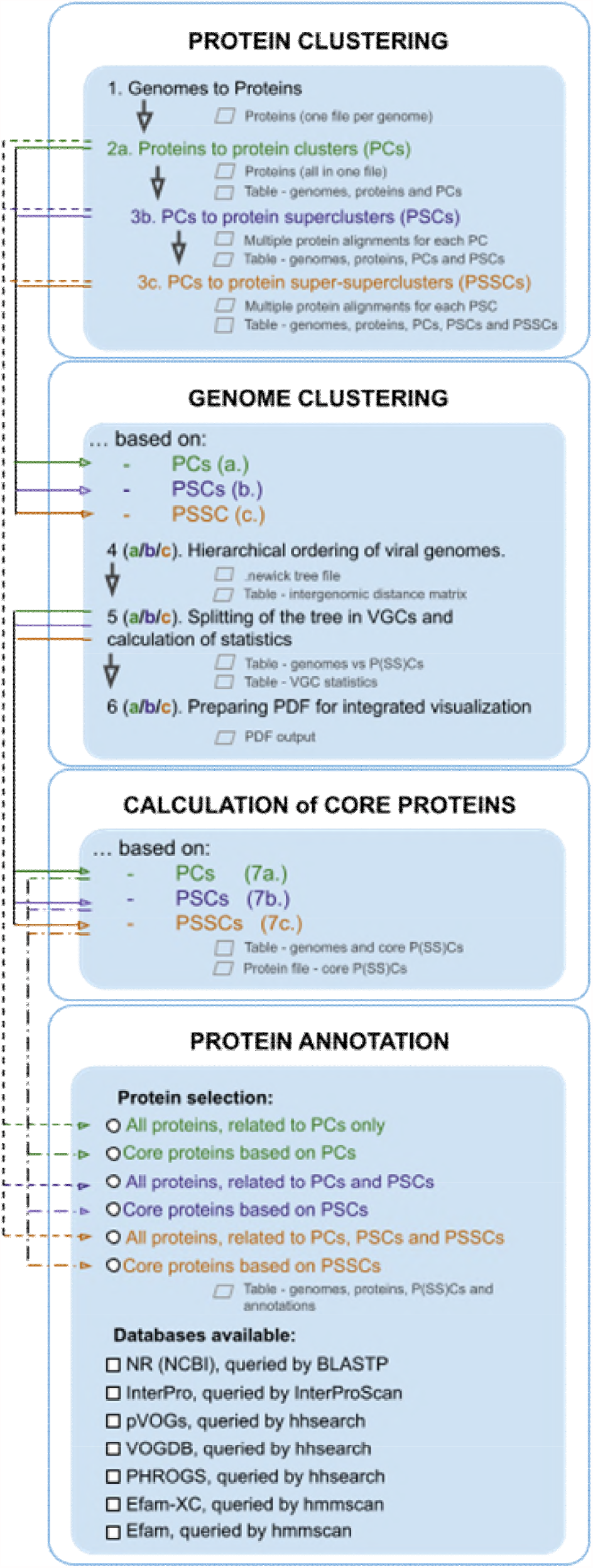
VirClust – module organization and possible workflows. The workflows on brach a are color coded green, on branch b are color coded purple and on branch c are color coded orange. The ussable outputs are mentioned for each step.

#### Step 1 – Protein prediction

In a first step, VirClust uses MetaGeneAnnotator (Noguchi *et al*. 2008) to predict genes in each viral genome, and then the seqinr R package (D. Charif and J.R. Lobry. 2007) to translate the predicted genes. Usable outputs from this step are the protein files, one .faa file per genome.

#### Step 2a – From proteins to protein clusters

In a second step, VirClust groups similar proteins into protein clusters (PCs). First, it compares all proteins with each other using BLASTP from the BLAST+ package (Camacho *et al*. 2009). The BLASTP hits are filtered based on their e-value, bitscore and coverage (of both subject and query). The default filtering parameters are bitscore > 50, e-value < 0.00001 and coverage = 0. Further, the remaining hits are used to cluster the proteins based on their i) e-values, ii) log10 transformed e-values, capped at 200 (the default) iii) bitscore or iv) normalized bitscores (maximum from “bitscore for prot1-prot2 hit / bitscore for prot1-prot1 hit” and “bitscore for prot2-prot1 hit / bitscore for prot2-prot2 hit”). The clustering is performed with with mcl (https://micans.org/mcl/), options “-I 2 --abc –o”. Usable outputs from this step are: i) one .faa file with the proteins from all genomes, labeled using a protein ID unique for the project; and ii) one .tsv file with all genes from all viral genomes (one gene per row), their genome location, their corresponding proteins (including the unique protein IDs) and the assigned PCs.

#### Step 3b – From protein clusters to protein superclusters

In a third, optional step, VirClust groups the PCs and their corresponding proteins into protein superclusters (PSCs), based on HMM similarities. First, for each PC calculated above it creates a multiple alignment with Clustal-Omega (Sievers and Higgins 2018), options “--pileup --iter=2”. Then it calculates Hidden Markov Models (HMMs) with hhmake (hhsuite package (Remmert *et al*. 2011), options “-id 100 -diff 1000000”). Further, it compares all HMMs with each other using hhsearch (hhsuite package, options “-id 100 -diff 0 -p 50 -z 1 -Z”). The results of this comparison can be filtered based on probability, coverage and alignment length, with thresholds established by the user. The default thresholds for filtering were those previously used for organizing dsDNA viral genomes into a bipartite network (Iranzo *et al*. 2016): i) probability >= 90, subject coverage >= 50 and ii) probability >= 99, subject coverage >= 20, alignment length >= 100. Finally, the hits passing the thresholds are used to cluster the PCs into PSCs, using mcl (options “-I 2 -te 20 -o”). The clustering can be done either based on e-values or on log10 transformed e-values (default). The usable outputs from this step are i) multiple alignments corresponding to each PC, in aligned multifasta format; and ii) a .tsv file, containing all genes from all viral genomes (one gene per row), their genome location, their corresponding proteins (including the unique protein IDs) and the assigned PCs and PSCs.

#### Step 3c – From protein superclusters to protein super-superclusters

In this optional step, VirClust groups PSCs into protein super-super clusters (PSSCs). For this, after creating a protein multiple alignment for each PSC, it proceeds similarly as at step 3b. The usable outputs from this step are i) multiple alignments corresponding to each PSC, in aligned multifasta format; and ii) a .tsv file, containing all genes from all viral genomes (one gene per row), their genome location, their corresponding proteins (including the unique protein IDs) and the assigned PCs, PSCs and PSSCs. From here on, the term P(SS)C is going to be used when referring generally to clusters of proteins, instead of using the longer of “PC, PSC or PSSC”.

#### Step 4a, 4b and 4c – Hierarchical clustering of the viral genomes bases on their PC, PSC or PSSC content

In this step, VirClust first calculates pairwise intergenomic distances, based either on the PC (4a), or on PSC (4b) or on PSSC (4c) content of each viral genome. For this, the presence of a P(SS)C in a viral genome is rewarded a score of 1, irrespectively of how many P(SS) replicates are found in the genome, and the absence of a P(SS)C is rewarded a score of 0. Pairwise distances are calculated using the formula:

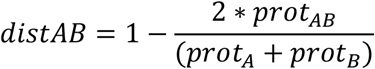

where,

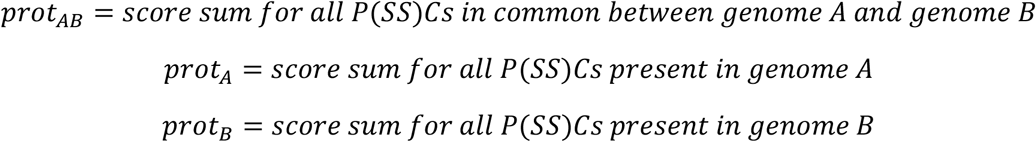

Further, VirClust performs a hierarchical clustering of the viral genomes based on the above-described intergenomic distances. For clustering it uses either the stats 3.5 package, without bootstrapping (default), or the pvclust 2.2 package (Suzuki and Shimodaira 2006; Shimodaira and Terada 2019), when bootstrap resampling is desired. The “complete” agglomeration method is used as default, with the other option being “average”. Following bootstrap resampling, the pvclust package calculates and reports three probability values for each cluster: i) selective inference p-value (SI); ii) approximately unbiased p-value (AU) and iii) bootstrap probability (BP) value (Shimodaira and Terada 2019). Due to the high CPU demand, the boot-strap option is inactivated if more than 50 genomes are inputted in VirClust web service, but is fully available in the stand-alone version. Usable outputs from this step are: i) a .tsv file, containing the calculated intergenomic distances; ii) a .newick file, containing the clustering results (the hierarchical tree). If bootstrapping is performed, then three .newick files are generated, one each for the SI, AU or BP values.

#### Step 5a, 5b and 5c – Splitting into viral genome clusters and related statistics

In this step, VirClust can split the viral genomes into clusters, by “cutting” the tree calculated in step4 at user defined distances. This tree cutting is performed with the “stats” package from R. The resulting viral genome clusters (VGCs) will contain viruses that are more similar to each other than to other viruses, either based on PC (5a), PSC (5b) or PSSC (5c) content.

Then, for each viral genome, VirClust calculates the following statistics: i) total number of proteins in the genome (corresponds to the total gene number); ii) total number of proteins that belong to singletons (P(SS)Cs containing only one protein, that is P(SS)Cs which are not shared with any other virus in the dataset); iii) total number of proteins found in P(SS)Cs shared with other viral genomes in the dataset; iv) total number of proteins found in P(SS)Cs shared with viral genomes from the same VGC, regardless if they are shared with viral genomes from outside the VGC as well; v) total number of proteins found in P(SS)Cs shared exclusively with viral genomes only from the same VGC; vi) total number of proteins found in P(SS)Cs shared with viral genomes from other VGC, regardless if they are shared with viral genomes from the same VGC as well; vi) total number of proteins found in P(SS)Cs shared exclusively with viral genomes from other VGC; vii) silhouette width (calculated with the R package “cluster” (Martin Maechler *et al*. 2021)).

Finally, a table is prepared in which the rows represent the viral genomes, ordered as in the tree calculated at step 4, and the columns represent the shared P(SS)Cs. The column order is based on their clustering with the “stats” R package, using the “binary” distance and the “complete” agglomeration method.

Usable outputs from this step are: i) a .tsv file with the genome and VGC statistics; and ii) a .tsv file with which P(SS)Cs are found in which viral genome, ordered as above.

#### Step 6a, 6b and 6c – Integrated visualization of the viral hierarchical clustering and of the protein content distribution

In the 6^th^ step, VirClust uses the R package ComplexHeatmap v. 2.5.3 (Gu *et al*. 2016) to generate a visual representation of the genome clustering. This is composed of: i) a clustering tree, as generated at step 4; ii) a heatmap documenting the presence of the different PCs (6a), PSCs (6b) or PSSCs (6c) in the viral genomes; iii) several annotations documenting the genome and protein statistics generated at step5, the silhouette width and the cluster designation. If the genomes have been split into several clusters, the heatmap and the corresponding annotation are split as well. The usable output from this step is a .PDF file with the visual representation.

#### Step 7a, 7b and 7c – Calculation of ore proteins

For each of the VGCs generated at step 5, VirClust is calculating the core PCs (7a), PSCs (7b) or PSSCs (7c), defined as P(SS)Cs found in all viruses from the respective VGC. For each VGC, the following usable outputs are generated: i) a .tsv file with the corresponding core proteins and their features (genome location, length, etc); and ii) a .faa with all corresponding core proteins.

#### Step 8 – Protein annotation

In a final step, VirClust is assisting the user to annotate the viral proteins. Homologues of the viral proteins are searched in several databases, as follows.

The NR database from NCBI is searched using BLASTP (“-evalue 0.0001 –max_target_ses 1000”). From the results, hits are removed if they represent hypothetical proteins, have a query/subject coverage < 40, have a pident < 30 or a bitscore < 50. From the remaining hits, that with the higher bitscore is used to annotate the query protein.

The prokaryotic Virus Orthologous Groups (pVOGs) database (Grazziotin *et al*. 2017), the Virus Orthologous Group database (VOGDB, https://vogdb.csb.univie.ac.at, (Kiening *et al*. 2019)) and the Prokaryotic virus Remote Homologous Groups (PHROGS) database (https://phrogs.lmge.uca.fr/index.php) are searched using hhsearch (Steinegger *et al*. 2019) (“-id 100 - diff 0 -p 50 -z 1 -Z 600”). Only hits with an e-value lower than 0.01 are kept. For each database, the hit with the highest score is used to annotate the query protein.

The efam and efam_XC databases are searched using hmmscan (Eddy 2011), options “-E 0.01”, followed by result removal if score < 40. For each database, the hit with the highest score is used to annotate the query protein.

The InterPro database (Finn *et al*. 2017) is searched using InterProScan (Jones *et al*. 2014). Results with the description “Domain of unknown function” and IP analysis “MobiDBLite” are removed.

The annotation results from all searched databases, including correspondig e-values and scores, are merged to the table containing the genome/protein information. The InterPro results are available both as a separate table or as a merged table. The tables can be downloaded in .tsv format.

### Running VirClust on a test dataset

For testing VirClust, a dataset of 942 dsDNA viral genomes (dataset A) was selected from all viral taxons currently recognized by the International Committee on Taxonomy of Viruses (ICTV), as found in the ICTV Master Species List 2020.v1 (https://talk.ictvonline.org/files/master-species-lists/m/msl/12314). The dataset included viruses from three viral realms. From *Duplodnaviria*, the following *Caudovirales* families were selected: *Herelleviridae, Demerecviridae, Autographiviridae, Ackermannviridae, Drexlerviridae, Chaseviridae, Salasmaviridae, Rountreeviridae, Schitoviridae, Zobellviridae* and *Guelinviridae*. From *Adnaviria*, all representatives were selected. From *Varidnaviria*, the following *Tectiliviricetes* were selected: *Kalamavirales* and *Vinavirales*, and *Autolykiviridae*.

From this dataset, a second, smaller dataset (dataset B) containing all ICTV recognized members of *Chaseviridae, Rountreeviridae* and *Zobellviridae* was assembled, for the illustration of the different VirClust features.

## Results and discussions

### VirClust – a tool for viral genome clustering, core protein detection and protein annotation

VirClust is a multifaceted viral genome analysis tool, developed to assist in the taxonomical classification of prokaryotic viruses and functional annotation of their protein encoding genes. To enable viral classification, on one hand side it performs a hierarchical clustering of the viral genomes, which can be used to group viruses at different taxonomic levels, and on the other hand side it identifies core proteins, which can be used for further phylogenetic analysis. To enable protein annotation, VirClust searches for homologous proteins within seven different protein sequence and HMM profiles databases.

VirClust is organized into four modules (see Figure 1): i) protein clustering; ii) genome clustering; iii) calculation of core proteins and iv) protein annotation.

In the first module, VirClust performs a series of basic steps (see Figure 1): 1) protein prediction and translation; 2a) protein grouping in PCs, based on BLASTP detectable homologies; 3b) PC grouping in PSCs, based on HMM profile search detectable homologies; and 3c) PSC grouping in PSSCs, again based on HMM profiles. The use of HMM profiles will enable the detection of more distantly protein homologies. The grouping of proteins into PSSCs is based on HMM profiles constructed from more diverrgent proteins sequences (from the PSCs). This procedure, of grouping PSCs into PSSCs has not been thoroughly explored evaluated by the scientific community. Therefore, the PSSC-based results should be considered with care.

In the next modules, VirClust can follow three parallel braches (see Figure 1, green, purple and orange arrows). All branches have identical steps, but the first (a) is based on PCs, the second (b) is based on PSCs and the last (c) is based on PSSCs (see Figure 1).

In the genome clustering module, VirClust uses the presence/absence of P(SS)Cs in viral genomes to calculate intergenomic distances. These distances are further used to cluster viruses hierarchically (step 4) and then to split them into VGCs based on a user defined distance threshold (step 5). Depending on the distance threshold used, the VGCs can be assigned to different taxonomic ranks (see below for a discussion).

Several indicators calculated in steps 4 and 5 serve to evaluate the clustering. For each individual cluster in the hierarchical tree, three different probability values (SI, AU and BP) are calculated by bootstrapping the P(S)Cs and can be used to assess the clustering uncertainty (Suzuki and Shimodaira 2006; Shimodaira and Terada 2019). In addition, the shared protein stats and the Silhouette width are genome specific statistics that can be used to appraise the affiliation of individual viruses to VGCs. The proportion of proteins shared with any other viral genomes in the analysed dataset shows what proportion of all proteins from a single virus are actually used for clustering. If only a small proportion of proteins are shared with other viruses in the dataset, it can increase the clustering uncertainty, because the singletons can hide relationships with yet unknown viruses and potentially, a different clustering. The Silhoutte width measures, on a scale of -1 to 1, how related is a virus with other viruses in the same VGCs. Values closer to 1 indicate higher similarity to members of its own VGC. Values closer to -1 indicate higher similarity with viruses in other VGCs. Similar to a negative Silhoutte width, a high proportion of proteins shared outside its own VGC can indicate an incorrect clustering.

A key feature of VirClust is the integrated visualization (step 6, see Figure 2) of the hierarchical clustering of viruses, of the distribution of their protein content and of their grouping in VGCs, with the corresponding statistics. The protein content of the viral genomes is visualized as a heatmap, in which the columns are representing PCs (step 6a), PSCs (6b) or PSSCs (step 6c), and the rows are representing the viral genomes, ordered according to the hierarchical clustering tree. Only shared proteins are depicted in the heatmap, the proportion of singletons being shown as an annotation along the heatmap (see Figure 2, “shared proteins” statistics). Together, the heatmap and the annotated statistics allow to open the “black box” of the tree: the user can visualize and thus identify which P(S)Cs have contributed to the hierarchical clustering, can identify which distance threshold is best for splitting the tree in VGCs, and also, can judge the quality of the clustering. Furthermore, the heatmap allows the identification of P(SS)Cs characteristic for certain viral groups and of potential gene duplication / gene split events (by the increased number of a P(SS)C in a viral genome).

**Figure 2:**
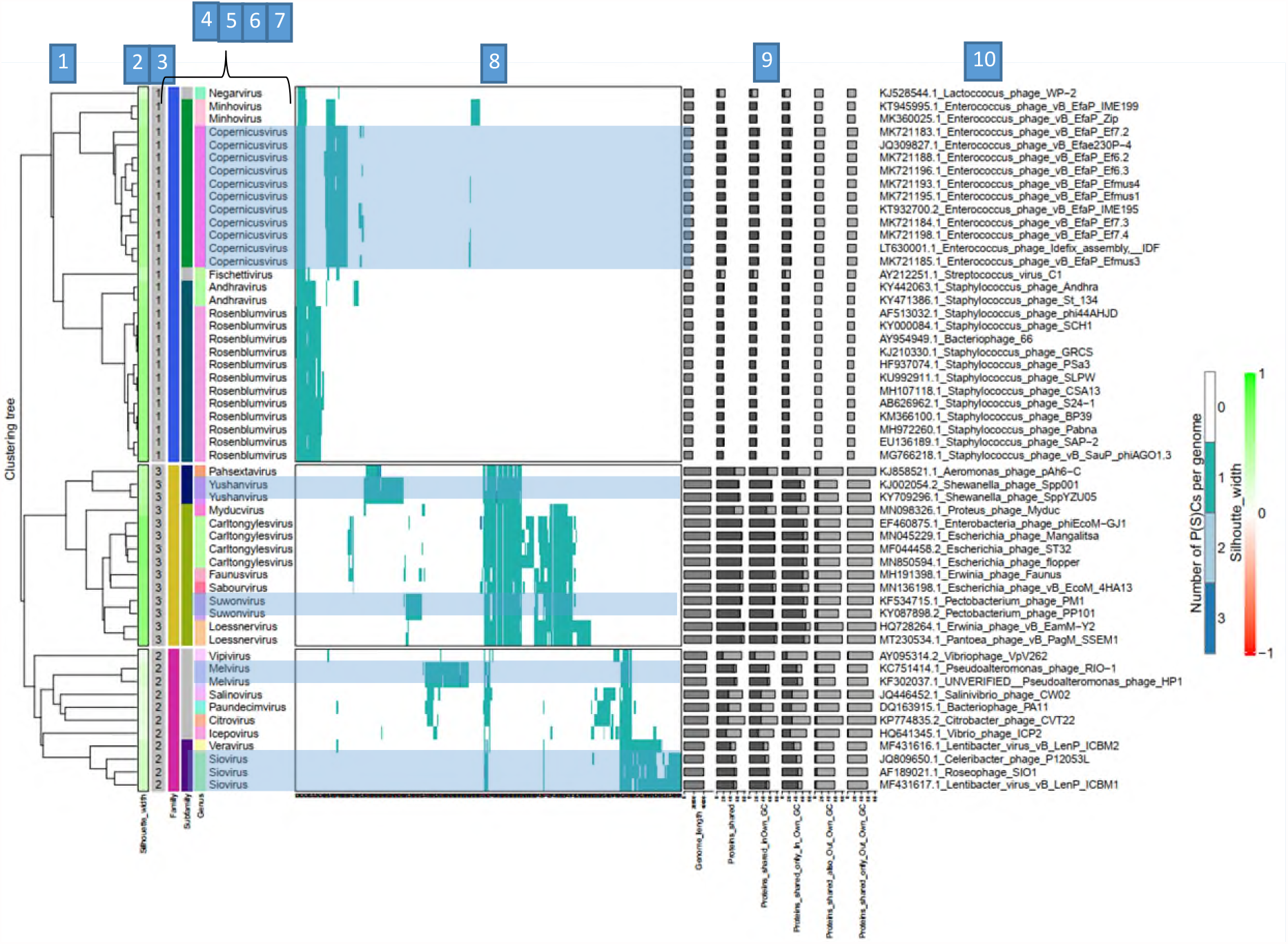
Integrated visualization of the viral clustering outputted by VirClust – example. A dataset comprised from all viral genomes in the *Zobellviridae, Chaseviridae* and *Rountreeviridae* was inputted in VirClust. The genome clustering was performed based on PCs with a bootstrapping of 1000 replicates. The resulting tree was split into VGCs using an 0.9 intergenomic distance threshold. The visual components are described further. **1. Hierarchical tree** calculated in step 4, using the PC based intergenomic distances. The stars represent > 95 % clustering p-values: blue – SI values; green – AU values. The clustering p-values depicted here have been manually retrieved from the .newick files generated at step 4, because they are not outputted in the integrated visualization. **2. Silhoutte width**, colour coded in a range from 1- (red) to 1 (green). **3. VGC ID**, as outputted in the genome statistic table from step 5. **The columns 4, 5, 6 and 7** are not part from the integrated visualization outputted by VirClust. They encode the official ICTV taxonomy of each viral genome analysed here. The taxonomic ranks encoded are: **4 – Family**; **5 – Subfamily; 6 and 7 – Genus. 8. Heatmap representation of the PC distribution in the viral genomes**. Rows are represented by individual viral genomes. Columns are represented by individual PCs. The ID of each PC can be read the bottom of the heatmap at image magnification. Colours encode the number of each PC per genome, with white signifying the PC absence, and the other colours signifying various degree of replication (from 1 to n, see legend). The blue-grey rectangles overlapping the heatmap and genus names are not part from the integrated visualization outputted by VirClust. They have been manually drawn here to illustrate how viruses in the same genus cluster together and have similar PC patterns, often distinct from the PC patterns of other viruses, including in the same VGC. **9. Viral genome specific statistics:** genome length, proportion of proteins shared (dark grey) from all proteins (light grey bar), proportion of proteins shared in own VGC, proportion of protein shared only in own VGC, proportion of proteins shared also outside own VGC and proportion of proteins shared only outside own VGC. **10. Virus name** (here including the GenBank accession number as suffix).

In the third module, VirClust calculates the corresponding core proteins for each of the VGCs identified at step 5. The core proteins are defined as those P(SS)Cs present in all the genomes from a VGC, regardless of their copy number per genome. The suitability of each identified core P(SS)C to be further used for phylogenetic analyses should be judged by the user from their multiple alignments (provided for download at steps 3b and 3c) and from their functional annotations (provided at step 8). P(SS)C subjected to gene duplication events, which can lead to truncated proteins, as well as those having gene insertions (as for example homing endonucleases, commonly spread in polymerases for example) should be carefully evaluated. Furthermore, proteins composed of multiple domains (for example DNA polymerases) can be encoded by a single gene or by more genes, each for a single domain. Depending on the stringency of the filtration thresholds at the protein clustering steps, the genes for the multiple domains can be grouped in a single P(SS)C (see more in section “Protein clustering – parameters choice”). The use of these P(SS)Cs for phylogenetic analysis should be carefully evaluated and eventually, the single domain proteins concatenated.

In the last module, VirClust performs protein annotations of the selected proteins (either all proteins, or only the core P(SS)Cs) by comparison with several sequence and HMM profiles databases (see Figure 1). Each database can be queried separately. The best hits from each database are identified for each protein. The results are then integrated in a single table, together with the information about the genome localization of each protein, and the assignment to P(SS)Cs and VGCs. The final annotation, integrating the information from all queried databases should be decided by the user, during careful evaluation of the annotation table. The protein assignment to P(SS)Cs greatly facilitates the annotation of those proteins without significant hits with any databases, because proteins grouped in the same P(SS)C should in general have the same function. The exceptions are those proteins composed from multiple domains or those with insertions. Evaluation of the annotation results and of the multiple alignments enable the identification of multiple domains, as well as of potential gene insertions.

The data analysis is organized by VirClust into projects. A project is defined by the project folder and the input genomes. Within a project, the complete workflow can be ran at once or in a stepwise manner. Furthermore, provided that they are not dependent on the results of previous steps, individual steps can be ran separately (see Figure 1). With the exception of step 1, all the other steps can be repeated with different parameters. If a step has been repeated, all the previously-calculated results from the next co-dependent steps are deleted, to avoid confusions. For example, the user can run only the first 2 steps (protein prediction and protein clustering) and return later to perform the genome clustering and core protein calculation based on PCs (see Figure 1, full green arrows). Initially, the user can chose to perform hierarchical ordering with no bootstrapping (step 4a) and to split the viruses into VGC with a distance threshold of 0.90 (step 5a). Upon inspection of the integrated visualization PDF (generated at step 6a), the user decides to try a distance threshold of 0.75. In this case, steps 5a, 6a and 7a have to be recalculated.

## Availability

The VirClust web-service (virclust.icbm.de) provides a graphical interface for running VirClust remotely. To avoid a heavy burden on the hosting server, it should be used only for small and medium sized projects. Larger projects can be analysed with VirClust stand-alone, which can be installed on user’s own servers, can be run from the command line and integrated into bioinformatics pipelines. Several operations are computationally intensive and have been parallelized: i) the BLASTP in step 2; ii) the HMM profile search in step3; iii) the bootstrapping in step 4 and v) the search in different databases in step 8. The computational time increases with the number of proteins / P(SS)Cs, especially during bootstrapping.

### Protein clustering – parameters choice

The protein clustering into PCs and further into PSCs and PSSCs represents the foundation on which the clustering of the viral genomes is based. It is a two-step process, in which first homologues are detected based either on BLASTP (for PCs) or HMM searches (for PSC, PSSC), and then the proteins are clustered based on the found homologies. Therefore, the parameters for defining homologues will influence which proteins cluster together. A literature review showed that, when determining PCs, in the first step the search results can be filtered based on their e-value, bitscore and alignment coverage of the two sequences. Different studies have used different combinations of the three parameters (for example, only e-value and bitscore (Roux *et al*. 2015)), or e-value and coverage (Zayed *et al*. 2021), or different values of the parameters, or even, no filtering at all (Enright *et al*. 2002). In the second step, the clustering can be based on e-values (Roux *et al*. 2015), log trasformed e-values (Enright *et al*. 2002) or normalized bitscore (Chan *et al*. 2013). When determining P(S)SCs, the results can be filtered based on their probability, coverage and HMM length (Iranzo *et al*. 2016). To enable detection of more distant homologues, Iranzo et al, 2016, have performed a two tier filtering, selecting all hits with i) a probability higher than 90% and coverage >50% and ii) a probability higher than 99%, but a coverage > 20% and minimum length of 100. VirClust uses all these parameters for protein clustering at PC, PSC and PSSC level, including the two tier filtering step for HMM results. However, rather than imposing strict values for these parameters, VirClust gives the user the opportunity to set her/his own values, in addition to the default suggestions.

There are viral proteins composed of multiple domains, which in some viruses can be encoded by the same gene, and in others separately. This is the case of the DNA polymerases from *Zobellviridae*, and we used this dataset to determine how the different filtering parameters can influence clustering of such proteins. Most genomes in this family have a DNA polymerase gene with an exonuclease and a polymerase domain. However, a few of them have the two domains as independent genes. From the three hit filtering parameters, the coverage parameter will most likely influence how these proteins will be clustered. When clustered with the default settings (for PCs – coverage > 0%; for PSCs – coverage 1 > 50% and coverage 2 > 20%), in which the coverage does not play a significant role, all DNA-pol related proteins (having both domains, or just the exonuclease domain, or just the polymerase domain) clustered in a single PSC. When increasing the coverage threshold (for PCs: coverage > 70%; for PSCs – coverage > 60%), the exonuclease and the polymerase domains clustered in different PSCs. However, also the DNA polymerase domains were split into three PSCs, indicating that recognition of more distantly related homologues was hampered by the increased coverage threshold (see SI file 3), even when using the more sensitive HMM searches. Depending on the purpose of the analysis, the user can set more relaxed thresholds, being aware that some clumping might occur, or more stringent thresholds, if a finer resolution of the protein clusters is needed.

### VirClust hierarchical clustering matches ICTV virus classification

A dataset of 942 dsDNA viruses was used to test VirClust’s ability to capture relationships between viruses at different taxonomic levels, when using PC and PSC based intergenomic distances. This dataset was assembled from ICTV recognized representatives of 3 viral realms and 16 families. The hierarchical trees produced in steps 4a and 4b were compared with the current ICTV taxonomy of the respective viruses (see Figure 3 and SI files 1 and 2).

**Figure 3:**
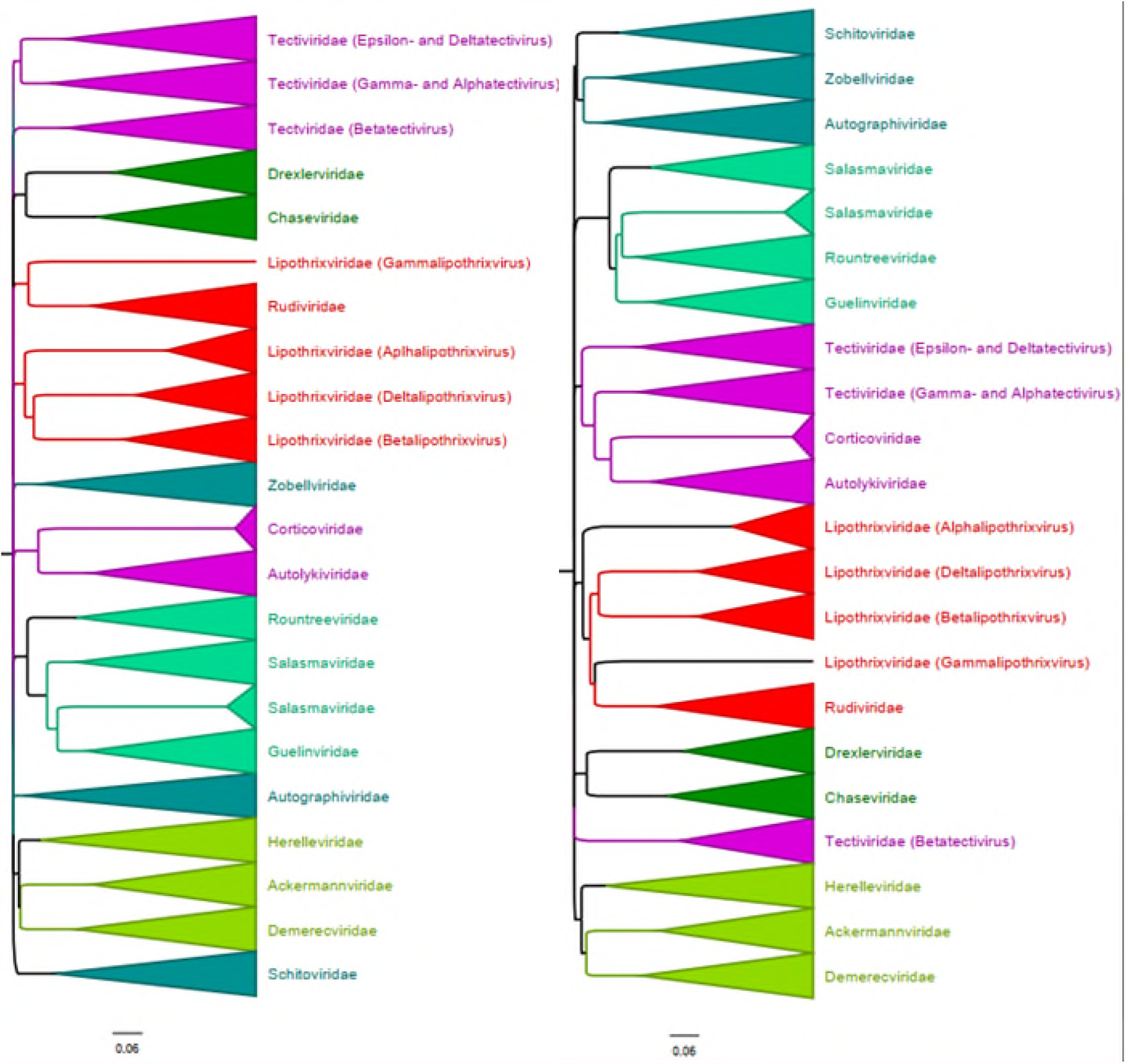
Hierarchical trees produced by VirClust based on PCs (a) and PSCs (b) for the large dsDNA viruses dataset. The newick files outputted by VirClust have been imported in FigTree v. 1.4.4, larger clusters collapsed and annotated. In red – clusters belonging to Adnaviria. In purple – clusters belonging to Varidnaviria. In different shades of green – clusters belonging to Duplodnaviria.

Generally, viral families formed major clusters, often branching directly from the root. No clustering at the realm level was evident in either of the two trees. This signifies that intergenomic distances based on PCs or PSCs are not able to capture distant relationships between all members of a viral realm. The exception were the viruses in the *Adnaviria* realm, which formed a single cluster in the PSC tree (see Figure 3B). However, this realm has only few members. Within *Duplodnaviria*, each family formed its own cluster. In both the PC and PSCs trees, several families grouped together into larger clusters, potentially representing order level taxons: i) *Drexlerviridae* and *Chaseviridae*; ii) *Rountreeviridae, Salasmaviridae* and *Guelinviridae*; iii) *Herelleviridae, Ackermannviridae* and *Demerecviridae*. In the PSC tree, the *Autographiviridae, Zobellviridae* and *Schitoviridae* formed a single cluster, relationships not captured in the PC tree.

Within *Varidnaviria* and *Adnaviria*, the *Tectiviridae* and the *Lipothrixviridae* families did not form single clusters. The betatectiviruses clustered always separately from the other tectiviridae. Even in the PSC tree, where most *Tectiviridae* clustered with *Corticoviridae* and *Autolykiviridae*, the other two *Varidinaviria* families, the *Betatectiviruses* clustered independently. This indicates that the taxonomic placement of betatectiviruses might need to be re-evaluated. The single gammalipothrixvirus clustered with the *Rudiviridae*, instead of the other lipothrixviridae, again indicating a need to re-evaluate its taxonomic placement.

Within the family-level clusters, both in the PC and PSC trees, the viral genomes were organized into sub-clusters in agreement with their taxonomic assignment at subfamily and genus level (see SI Files 1 and 2). Together, these data show that the hierarchical clustering produced by VirClust matches the current ICTV classification at family, subfamily and genus level.

Organizing viruses into hierarchical taxonomic ranks implies that some type of distance thresholds are assigned to each rank. With the exception of the species and the genus levels, these thresholds have not been yet explicitly defined by the Bacterial and Archaeal Viruses Committee of ICTV. Furthermore, it is not yet clear if the same thresholds could or should be used across all virus realms. However, it is clear that inside one realm, there is a need apply coherent thresholds, to ensure that the viral diversity is hierarchically distributed and comparable across taxons. Here, different distance thresholds (0.98, 0.90 and 0.85) were applied to the hierarchical trees produced by VirClust, to determine if one of them is suitable for defining family level clusters. The results indicate that different thresholds apply to the three realms (see Table 1), likely reflecting the variations in taxon-level specific classification criteria.

**Table 1:**
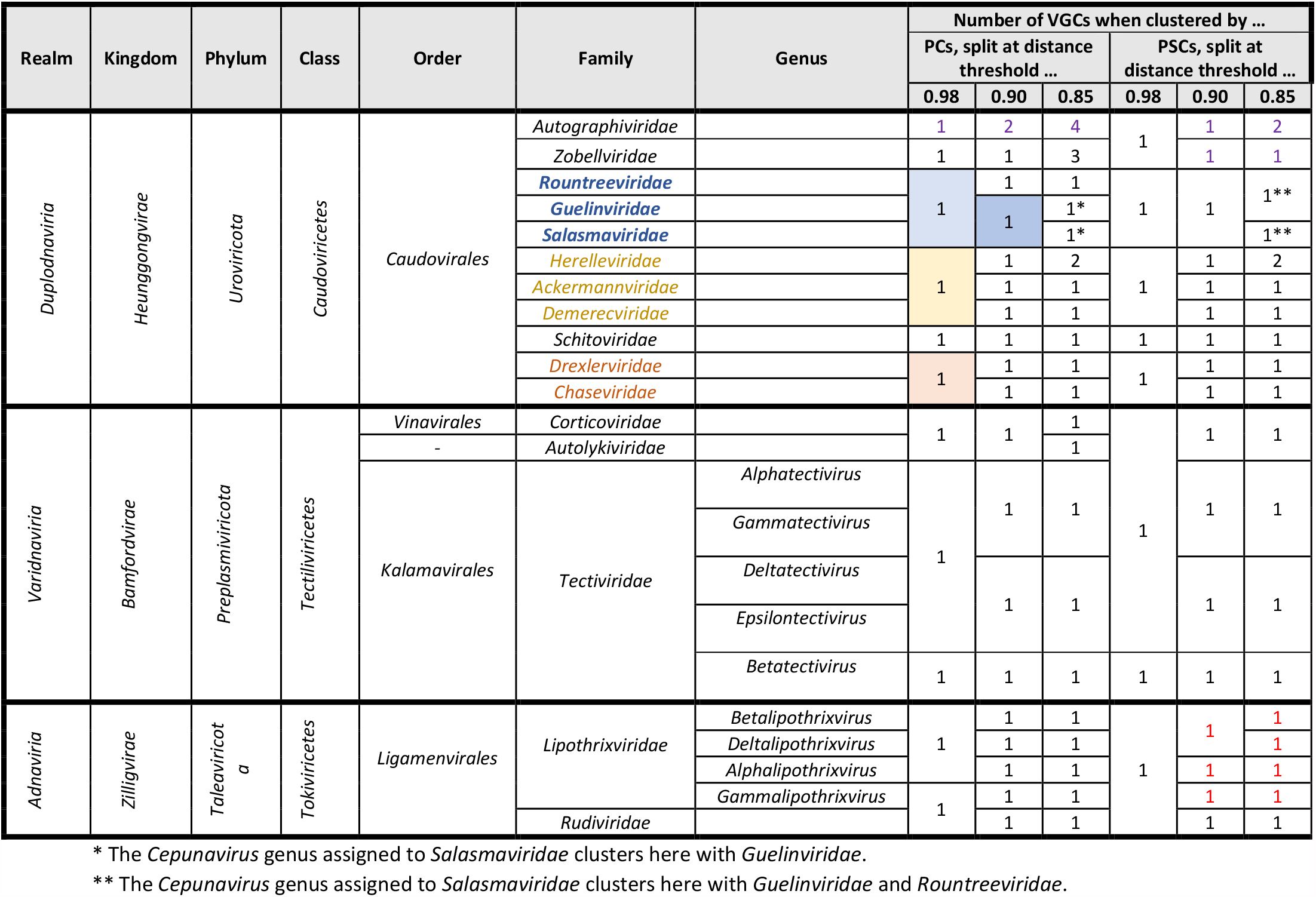
Spliliting of the PC and PSC trees into VGCs at different distance thresholds.

Within *Duplodnaviria*, a threshold of 0.90 intergenomic distance applied on the PC tree delineated most families, the other two thresholds resulting in either a higher clumping or higher splitting of the families. The exceptions were the *Autographiviridae* family, which at 0.90 was split in 2 VGC, and the *Guelinviridae* and *Salasmaviridae* families, which formed a single VGC. Potentially, 0.98 could be used for order level delineation.

For the other two realms, no clear-cut threshold delineating families was found, pointing toward a variability into the criteria used for delineation. Within *Varidnaviria*, a 0.85 distance could discriminate between the *Corticoviridae* and *Autolykiviridae* families, but only a 0.98 distance threshold brought together most of *tectiviridae* (with the exception of *Betatectiviruses*, which clustered separately). PC based intergenomic distances between viruses in the *Corticoviridae* and *Autolykiviridae* families were thus smaller than distances between viruses in the *Alphatectivirus* and *Deltatectivirus* genera of *Tectividae* (see SI File 1 and Table 1). This indicates a need to re-evaluate these taxons. Within *Adnaviria*, the same threshold that delineated the *Rudiviridae* family actually discriminated between individual genera of the *Lipothrixviridae* family. This indicates that intergenomic distances between viruses within the *Rudiviridae* family were similar to intergenomic distances between viruses within individual *Lipothrixvirus* genera (see also SI Files 1 and Table 1).

The thresholds identified here (for example 0.9 for family level within *Caudovirales*) apply only when protein clustering is performed with the default parameters of VirClust. Increasing the stringency criteria for protein cluster formation (for example by adding a filtering step based on coverage) will most likely increased the observed distances between the taxons and the thresholds should be re-evaluated.

VirClust produces viral genome clusters matching the current prokaryotic virus taxons, at different taxonomic levels, from genus to family. With the help of distance thresholds, as for example 0.9 for families in the *Caudovirales*, the VirClust trees can be split into viral genome clusters of different taxonomic ranks. Furthermore, the core proteins, defining each VGC, can be easily identified and annotated. VirClust represents a new tool for the classification of prokaryotic viruses, which not only groups viruses into hierarchical clusters, but also enables the identification of the genomic features responsible for the respective classification. And hopefully, it will play a role in the disentangling of the current paraphyletic *Podoviridae, Siphoviridae* and *Myoviridae* families.

## References

Aiewsakun, P., Adriaenssens, E.M., Lavigne, R., Kropinski, A.M., and Simmonds, P. (2018) Evaluation of the genomic diversity of viruses infecting bacteria, archaea and eukaryotes using a common bioinformatic platform: steps towards a unified taxonomy. Journal of General Virology, doi: 10.1099/jgv.0.001110.

Aiewsakun, P., and Simmonds, P. (2018) The genomic underpinnings of eukaryotic virus taxonomy: creating a sequence-based framework for family-level virus classification. Microbiome, doi: 10.1186/s40168-018-0422-7.

Bin Jang, H., Bolduc, B., Zablocki, O., Kuhn, J.H., Roux, S., Adriaenssens, E.M., et al. (2019) Taxonomic assignment of uncultivated prokaryotic virus genomes is enabled by gene-sharing networks. Nature Biotechnology, doi: 10.1038/s41587-019-0100-8.

Bolduc, B., Jang, H.B., Doulcier, G., You, Z.-Q., Roux, S., and Sullivan, M.B. (2017) vConTACT: an iVirus tool to classify double-stranded DNA viruses that infect Archaea and Bacteria. PeerJ, doi: 10.7717/peerj.3243.

Camacho, C., Coulouris, G., Avagyan, V., Ma, N., Papadopoulos, J., Bealer, K., and Madden, T.L. (2009) BLAST+: architecture and applications. BMC bioinformatics, doi: 10.1186/1471-2105-10-421.

Chan, C.X., Mahbob, M., and Ragan, M.A. (2013) Clustering evolving proteins into homologous families. BMC bioinformatics, doi: 10.1186/1471-2105-14-120.

D. Charif, and J.R. Lobry. (2007) SeqinR 1.0-2: a contributed package to the R project for statistical computing devoted to biological sequences retrieval and analysis. In Structural approaches to sequence evolution: Molecules, networks, populations. U. Bastolla, M. Porto, H.E. Roman, and M. Vendruscolo (eds). New York: Springer Verlag, pp. 207–232.

Eddy, S.R. (2011) Accelerated Profile HMM Searches. PLoS computational biology, doi: 10.1371/journal.pcbi.1002195.

Enright, A.J., van Dongen, S., and Ouzounis, C.A. (2002) An efficient algorithm for large-scale detection of protein families. Nucleic Acids Research, doi: 10.1093/nar/30.7.1575.

Finn, R.D., Attwood, T.K., Babbitt, P.C., Bateman, A., Bork, P., Bridge, A.J., et al. (2017) InterPro in 2017-beyond protein family and domain annotations. Nucleic acids research, doi: 10.1093/nar/gkw1107.

Gorbalenya, A.E., Krupovic, M., Mushegian, A., Kropinski, A.M., Siddell, S.G., Varsani, A., et al. (2020) The new scope of virus taxonomy: partitioning the virosphere into 15 hierarchical ranks. Nature Microbiology, doi: 10.1038/s41564-020-0709-x.

Grazziotin, A.L., Koonin, E.V., and Kristensen, D.M. (2017) Prokaryotic virus orthologous groups (pVOGs). A resource for comparative genomics and protein family annotation. Nucleic acids research, doi: 10.1093/nar/gkw975.

Gu, Z., Eils, R., and Schlesner, M. (2016) Complex heatmaps reveal patterns and correlations in multidimensional genomic data. Bioinformatics (Oxford, England), doi: 10.1093/bioinformatics/btw313.

Iranzo, J., Krupovic, M., and Koonin, E.V. (2016) The Double-Stranded DNA Virosphere as a Modular Hierarchical Network of Gene Sharing. mBio, doi: 10.1128/mBio.00978-16.

Jones, P., Binns, D., Chang, H.-Y., Fraser, M., Li, W., McAnulla, C., et al. (2014) InterProScan 5: genome-scale protein function classification. Bioinformatics (Oxford, England), doi: 10.1093/bioinformatics/btu031.

Kazlauskas, D., Varsani, A., Koonin, E.V., and Krupovic, M. (2019) Multiple origins of prokaryotic and eukaryotic single-stranded DNA viruses from bacterial and archaeal plasmids. Nature communications, doi: 10.1038/s41467-019-11433-0.

Kiening, M., Ochsenreiter, R., Hellinger, H.-J., Rattei, T., Hofacker, I., and Frishman, D. (2019) Conserved Secondary Structures in Viral mRNAs. Viruses, doi: 10.3390/v11050401.

Koonin, E.V., Dolja, V.V., Krupovic, M., Varsani, A., Wolf, Y.I., Yutin, N., Zerbini, F.M., and Kuhn, J.H. (2020) Global Organization and Proposed Megataxonomy of the Virus World. Microbiology and Molecular Biology Reviews, doi: 10.1128/MMBR.00061-19.

Koonin, E.V., Senkevich, T.G., and Dolja, V.V. (2006) The ancient Virus World and evolution of cells. Biology direct, doi: 10.1186/1745-6150-1-29.

Krupovic, M., Dolja, V.V., and Koonin, E.V. (2019) Origin of viruses: primordial replicators recruiting capsids from hosts. Nature reviews. Microbiology, doi: 10.1038/s41579-019-0205-6.

Krupovic, M., and Koonin, E.V. (2017) Multiple origins of viral capsid proteins from cellular ancestors. Proceedings of the National Academy of Sciences of the United States of America, doi: 10.1073/pnas.1621061114.

Krupovic, M., Kuhn, J.H., Wang, F., Baquero, D.P., Dolja, V.V., Egelman, E.H., Prangishvili, D., and Koonin, E.V. (2021) Adnaviria: a new realm for archaeal filamentous viruses with linear A-form double-stranded DNA genomes. Journal of Virology, doi: 10.1128/JVI.00673-21.

Martin Maechler, Peter Rousseeuw, Anja Struyf, Mia Hubert, and Kurt Hornik (2021) cluster: Cluster Analysis Basics and Extensions. [WWW document]. URL https://CRAN.R-project.org/package=cluster.

Meier-Kolthoff, J.P., and Göker, M. (2017) VICTOR: genome-based phylogeny and classification of prokaryotic viruses. Bioinformatics, doi: 10.1093/bioinformatics/btx440.

Moraru, C., Varsani, A., and Kropinski, A.M. (2020) VIRIDIC-A Novel Tool to Calculate the Intergenomic Similarities of Prokaryote-Infecting Viruses. Viruses, doi: 10.3390/v12111268.

Nishimura, Y., Yoshida, T., Kuronishi, M., Uehara, H., Ogata, H., and Goto, S. (2017) ViPTree: the viral proteomic tree server. Bioinformatics (Oxford, England), doi: 10.1093/bioinformatics/btx157.

Noguchi, H., Taniguchi, T., and Itoh, T. (2008) MetaGeneAnnotator: detecting species-specific patterns of ribosomal binding site for precise gene prediction in anonymous prokaryotic and phage genomes. DNA research : an international journal for rapid publication of reports on genes and genomes, doi: 10.1093/dnares/dsn027.

R Core Team (2018) R: A language and environment for statistical computing. [WWW document]. URL https://www.R-project.org/.

Remmert, M., Biegert, A., Hauser, A., and Söding, J. (2011) HHblits: lightning-fast iterative protein sequence searching by HMM-HMM alignment. Nature methods, doi: 10.1038/nmeth.1818.

Roux, S., Enault, F., Hurwitz, B.L., and Sullivan, M.B. (2015) VirSorter: mining viral signal from microbial genomic data. PeerJ, doi: 10.7717/peerj.985.

Shimodaira, H., and Terada, Y. (2019) Selective Inference for Testing Trees and Edges in Phylogenetics. Frontiers in Ecology and Evolution, doi: 10.3389/fevo.2019.00174.

Sievers, F., and Higgins, D.G. (2018) Clustal Omega for making accurate alignments of many protein sequences. Protein science : a publication of the Protein Society, doi: 10.1002/pro.3290.

Steinegger, M., Meier, M., Mirdita, M., Vöhringer, H., Haunsberger, S.J., and Söding, J. (2019) HH-suite3 for fast remote homology detection and deep protein annotation. BMC bioinformatics, doi: 10.1186/s12859-019-3019-7.

Suzuki, R., and Shimodaira, H. (2006) Pvclust: an R package for assessing the uncertainty in hierarchical clustering. Bioinformatics, doi: 10.1093/bioinformatics/btl117.

Turner, D., Kropinski, A.M., and Adriaenssens, E.M. (2021) A Roadmap for Genome-Based Phage Taxonomy. Viruses, doi: 10.3390/v13030506.

Zayed, A.A., Lücking, D., Mohssen, M., Cronin, D., Bolduc, B., Gregory, A.C., et al. (2021) efam: an expanded, metaproteome-supported HMM profile database of viral protein families. Bioinformatics.

